# Preconception stress in mice alters maternal behavioral, neural, and endocrine outcomes, and produces sex-specific disruptions in offspring

**DOI:** 10.64898/2026.09.29.755005

**Authors:** Marissa M. Nicodemus, Laiklyn A.M. Luther, Isabella M. Linton, Mostafa A. Foual, Kendra Y. Maynard, Kathleen E. Morrison

## Abstract

Disruptions to maternal behaviors that emerge during the peripartum period have been shown to have lasting physiological and behavioral consequences for offspring. In addition to effects on offspring, disrupted maternal behaviors have been associated with peripartum mood and anxiety disorders. Therefore, it is critical for the wellbeing of both mother and offspring to identify upstream risk factors for the disruption of maternal behavior. We have previously found that pubertal stress leads to a blunted hypothalamic-pituitary-adrenal (HPA) axis response in peripartum humans and mice. Given that dysfunction of the HPA axis is associated with both PMADs and disrupted maternal behavior, we examined how pubertal stress influenced maternal behavior throughout peripartum in mice using a pup retrieval test. We found that pubertal stress led to faster pup retrieval, but a less predictable pattern of maternal behavior, as well as a blunted HPA axis response during pregnancy. We also examined whether maternal pubertal stress influenced offspring behavior in adolescence and adulthood. We found sex-specific differences in risk-taking behavior during adolescence that are no longer present in adulthood. Finally, we used *in situ* hybridization to examine gene expression during pregnancy in the medial preoptic area of the hypothalamus, a region implicated in maternal behavior. Following pubertal stress, we found cell-type specific changes in the expression of genes necessary for the expression of maternal behavior. Collectively, these findings contribute to our understanding of upstream risk factors for peripartum disruptions to maternal behavior and the lasting effects of pubertal stress on both mother and offspring.

## Introduction

The peripartum period is characterized by physiological and behavioral changes that prepare individuals for the transition to motherhood. Maternal behaviors that ensure the well-being of offspring emerge during peripartum, including an increased sensitivity to infant cries, altered mother-infant gaze, and germ-aversion [1–6]. Disruptions to maternal behaviors in humans are associated with lasting effects on offspring, including disruptions to cognition, brain development, and physiology [7–9], and are associated with peripartum mood and anxiety disorders (PMADs) [10]. For mothers, PMADs are associated with impairments in daily functioning and increased risk for the recurrence of depression and development of chronic physical disease years after the peripartum period [11]. For offspring, PMADs are associated with an increased risk for delays in social, language, and motor development [12].

Given these lasting and serious consequences, it is important to identify upstream risk factors. In humans and nonhuman animals, early life stress prior to conception has been identified as a risk factor for the disruption of maternal behavior during pregnancy and postpartum [13–18]. Such changes in maternal behavior have been associated with lasting effects in the offspring of mothers that underwent early life stress [16,19]. Early life preconception stress is also known to disrupt the hypothalamic-pituitary-adrenal (HPA) axis, which orchestrates the neuroendocrine stress response in mammals [13,17,18,20,21]. Dysfunction of the HPA axis is a known endophenotype of mood and anxiety disorders [22–25], and HPA axis function is known to mediate the relationship between maternal early life experiences and maternal behavior in a mother-infant interaction task [15]. Collectively, this evidence suggests that disruption of HPA axis function following early life preconception stress may play an important role in the expression of maternal behavior, which, in turn, is a key mechanism for transmission of risk from mother to offspring.

The majority of the mechanistic work on early life preconception stress focuses on stress during infancy [13,14,17,18]. However, there is a body of evidence suggesting that that females may be particularly vulnerable to the effects of stress that occurs around the time of adolescence or puberty [26–29]. Therefore, administering stress during puberty may be well-suited to capture the entire risk profile in female subjects. However, the lasting effects of pubertal stress on adult maternal behavior and physiology have only recently emerged as an area of focus.

We have addressed such gaps using our well-established model, which was developed to address risk factors relevant to female life experiences. Peripartum humans and mice that underwent chronic stress during puberty showed a blunted glucocorticoid response to acute stress, humans showed significantly higher scores on a measure of postnatal depression, and mice showed disrupted transcriptional and epigenetic responses in the brain in response to becoming pregnant [20,21,30,31]. In our human studies, male infants of pubertally stressed mothers had a lower cortisol response to stress compared to female infants. Importantly, this sex difference was not present in the offspring of mothers who reported no pubertal adverse experiences [32]. Here, we address broader consequences for behavior during the peripartum period and potential consequences for offspring in mice.

Additionally, the neural mechanisms underlying the impact of early life preconception stress on later behavior and physiology are only beginning to be understood. The medial preoptic area of the hypothalamus (mPOA) drives the expression of maternal care and suppresses aggression toward pups [33–35], and as such undergoes reorganization during pregnancy to accommodate these changes. Few studies have focused on the impact of early life stress on the mPOA, but emerging findings identify differences in cell density and transcriptional changes following early life stress [36–39]. Furthermore, the effect of early life preconception stress on the mPOA during pregnancy is largely unexplored. In addition to its role in the performance of maternal care, the mPOA is considered part of the extended HPA axis and acts as an inhibitor of the glucocorticoid stress response [40–43]. Considering both the inhibitory influence of the mPOA over the PVN and the reorganization of the mPOA throughout the peripartum window, it is possible that this circuit plays a role in the blunted HPA axis response we have previously observed.

Here, we examined HPA axis function, maternal behavior, and offspring outcomes in a single set of animals, allowing for critical associations to be made across outcomes that are known to be individually influenced by early life preconception stress [13–18,20,21,32]. Additionally, we characterized the expression of genes both necessary for the expression of maternal behavior and involved in the inhibition of the stress response [40–42,44]. Together, these studies inform our understanding of the effects of early life stress in the behavior and physiology of both mother and offspring, as well as provide insight into potential molecular underpinnings of disruption to these outcomes following early life stress (**Figure 1**).

**Figure 1.**
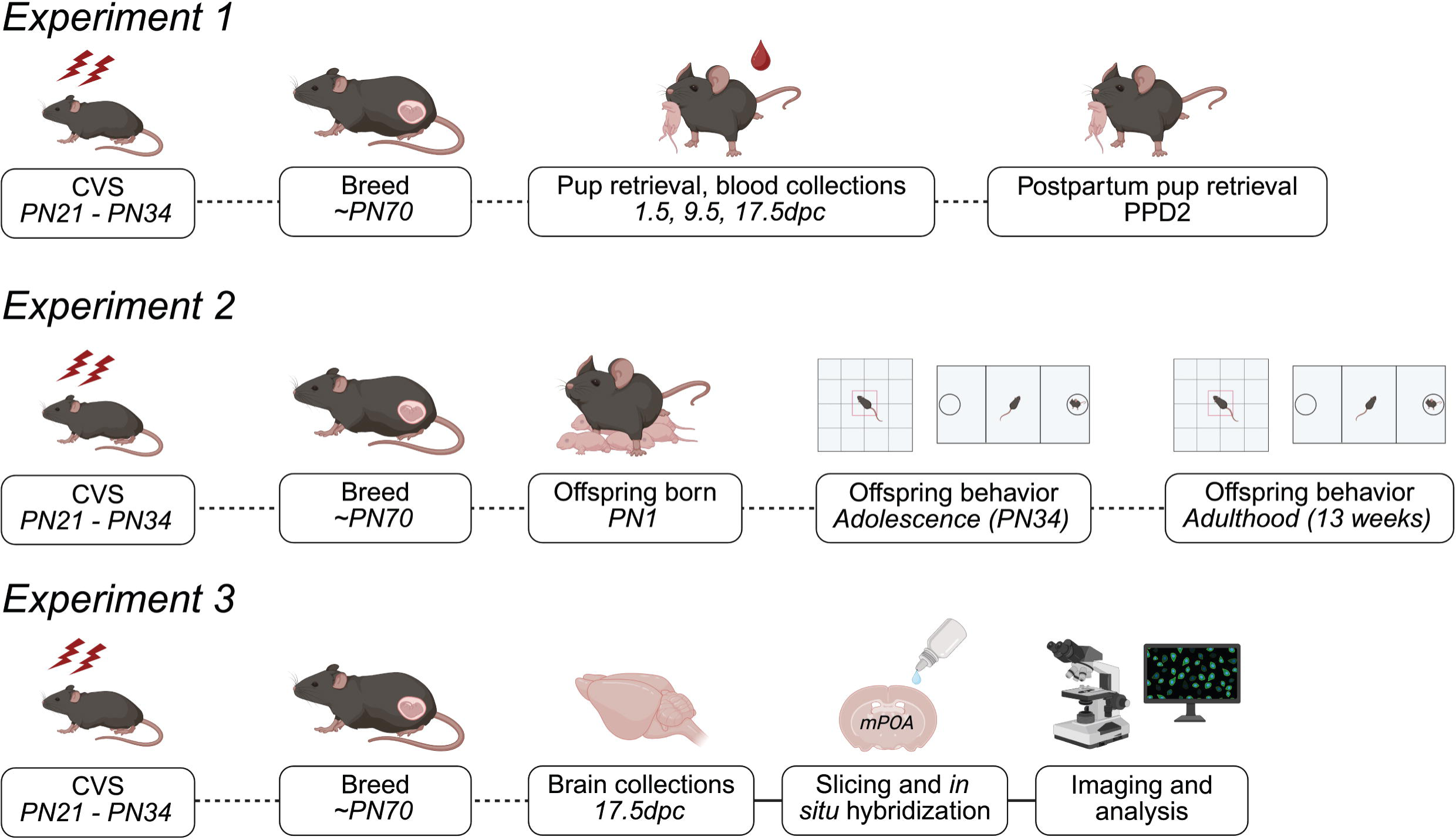
Timeline of experimental procedures. Animals underwent chronic variable stress (CVS) from postnatal days (PN) 21-34 or were left as Controls. All females were left undisturbed from PN35 until ten weeks of age, when they were bred to become pregnant. In Experiment 1, a pup retrieval test was administered at 1.5dpc, 9.5dpc, 17.5dpc, and PPD2. Following the tests during pregnancy, tail blood was collected to measure the corticosterone response. In Experiment 2, the offspring of dams from Experiment 1 underwent an open field test and a social preference test during adolescence and again in adulthood. In Experiment 3, brains were collected at 17.5dpc and the medial preoptic area (mPOA) was collected. RNAscope *in situ* hybridization, imaging, and quantification were performed. CVS = chronic variable stress; dpc = days post conception; PN = postnatal day; PPD = postpartum day. Created with BioRender.com.

## Materials and Methods

### General Methods

Extended details of the methods can be found in the Supplement.

#### Mice

Mice used were in-house, heterozygous Crh-IRES-Cre:Ai14, expressing tdTomato in CRF neurons on a mixed C57BL/6 x 129S1/SvImJ background (N = 40) or the offspring of such mice (N = 48). All procedures were approved by the West Virginia University Institutional Animal Care and Use Committee and were conducted in accordance with the National Institutes of Health Guide for the Care and Use of Laboratory Animals.

#### Pubertal Stress

Pubertal chronic variable stress (CVS) was conducted as previously [20,30,31]. Mice underwent 14 days of CVS starting on postnatal day (PN) 21, during which two stressors were administered simultaneously, once per day, for two hours: auditory stressors (owl screech, white noise), olfactory stressors (predator odor, ethanol odor), or tactile stressors (restraint in a conical tube, wet bedding, multiple cage change, exposure to novel stimulus, no bedding). See supplement for timing and stressor details (**Table S1**). CVS mice were singly housed at PN21 and for the duration of CVS, but were pair housed with same-sex, same-condition littermates at the end of stress. Control mice were weaned at PN28 and pair-housed with same-sex littermates as previously [20,21,30,31,45].

#### Breeding

At ten weeks old, female mice were bred with a male for 1-3 nights. If a copulation plug was observed during the first hour of the light cycle, the female mouse was considered 0.5 days post conception (dpc) and was singly housed in a clean cage.

### Experiment 1 - Maternal behavior and stress reactivity

#### Pup Retrieval

Adult female mice underwent a home cage pup retrieval task on four different days: 1.5 dpc, 9.5 dpc, 17.5 dpc, and postpartum day (PPD) 2. Testing began one hour after lights on. On days when test mice were pregnant, probe pups from a donor dam were used. One pup was placed in each of the corners far from the nest, with a male always placed on the left corner and a female on the right corner. The test female was returned to the nest, and behavior was recorded for 10 min. Tests ended early if mice acted aggressively toward pups. Behavior was scored in Anymaze (**Table S2**).

#### Glucocorticoid response to acute stress

On 1.5, 9.5, and 17.5 dpc testing days, 10 ul of blood was collected from females from a small nick at the end of the tail at 15, 30, and 120 minutes following the start of pup retrieval testing and processed as previously [21,31,45–54]. Corticosterone concentration was determined from blood plasma by ^125^I-corticosterone radioimmunoassay (MP Biomedicals) per manufacturer instructions.

### Experiment 2 - Offspring development and behavior

#### Offspring

One male and one female offspring from each dam from Experiment 1 was weaned at PN28 and pair-housed with a same-sex, same-maternal stress condition animal from a different dam. Offspring underwent Open Field and Social Preference testing during adolescence (PN34) and again in adulthood (13 weeks old). Tracking was performed by AnyMaze.

#### Open Field

Offspring were placed into the center of a 40 x 40 centimeter apparatus and allowed to freely explore for 5 minutes.

#### Social Preference

A 3-chamber box was used for social preference testing. Mice were allowed to habituate to the apparatus, which included two empty cups placed at the center of the far zones of the chamber, for 5 minutes. For the test, a same-sex, same-age novel probe mouse was placed into the cup in one of the far zones. Then, the test mouse was returned to the chamber and allowed to explore for 5 minutes. The social preference index was calculated as: (time in social zone - time in nonsocial zone)/(time in social zone + time in nonsocial zone).

### Experiment 3 - Hypothalamic expression of maternally relevant genes

#### Tissue collection and imaging

At baseline on the morning of 17.5 dpc, mice were anesthetized via 5% isoflurane, then euthanized via decapitation. Brains were extracted, immediately placed on dry ice, and stored at -80°C. Brains were cryosectioned at -20°C and the medial preoptic area (mPOA) was collected (0.25mm through -0.23mm relative to bregma). 4, 10um slices were collected in a set of three slides. Each slide in the set represented the full anterior to posterior range for the mPOA. Fluorescent images were acquired using a 40x objective on an Olympus BX43 Upright Microscope.

#### In situ hybridization

Gene expression analyses were performed using the RNAscope Fluorescent Multiplex Reagent Kit (version 2, ACDBio) according to manufacturer instructions for fresh frozen tissue (RNAscope Multiplex Fluorescent Reagent Kit v2 User Manual, 2022). Probes targeted *Esr1*, *Pgr*, and *Gal*. Gene expression was quantified in QuPath.

## Statistical Analyses

ANOVAs were used for most outcomes, with Tukey post-hoc testing as appropriate. For all ANOVAs, if the assumption of sphericity was violated, the Geisser-Greenhouse correction was used. Independent sample t-tests (Control vs CVS) were used to compare gene expression and cell population measures for *in situ* hybridization. A Markov Chain analysis was performed on the transition probabilities of maternal behavior during pup retrieval. The ROUT method was used to identify outliers. P value was set to 0.05. Full statistical outputs are presented in **Tables S3-S8**.

## Results

### Preconception pubertal stress led to changes in the organization of maternal care

We previously found that pubertal stress (CVS) interacts with pregnancy to lead to a blunted glucocorticoid response to an acute stressor. Here, we extended our findings to maternal behavior (**Figure 2A**). We found no effect of pubertal stress on dam body weight throughout pregnancy, the percentage of male pups in a litter, or the total number of pups in a litter (**Figure 2B-D**). Some mice were excluded from behavior analyses due to aggression toward pups during the test (n=4; 1 Control, 3 CVS; **Figure 2E**). Analysis of latency to first retrieval (**Figure 2F**) and time spent in the nest (**Figure 2G**) demonstrated that CVS females had a shorter latency to retrieve the first pup across all test days *(F_pubertal stress_*(1, 21) = 4.819, *p* = 0.04) and spent more time in the nest on 1.5dpc compared to Controls (*F_pubertal stress x test day_*(2.57, 53.89) = 2.96, *p* = 0.048; Tukey_1.5dpc_ *p* = 0.009). A more complex understanding of the organization of maternal behavior was examined using a Markov Chain Model to calculate transition probabilities between behaviors (**Figure 2H-I**). These transition probabilities revealed that the patterns of behavior during a pup retrieval task were altered by preconception pubertal stress a pattern during pregnancy where CVS mice were more likely to repeat the nest entry behavior before moving to another behavior.

**Figure 2.**
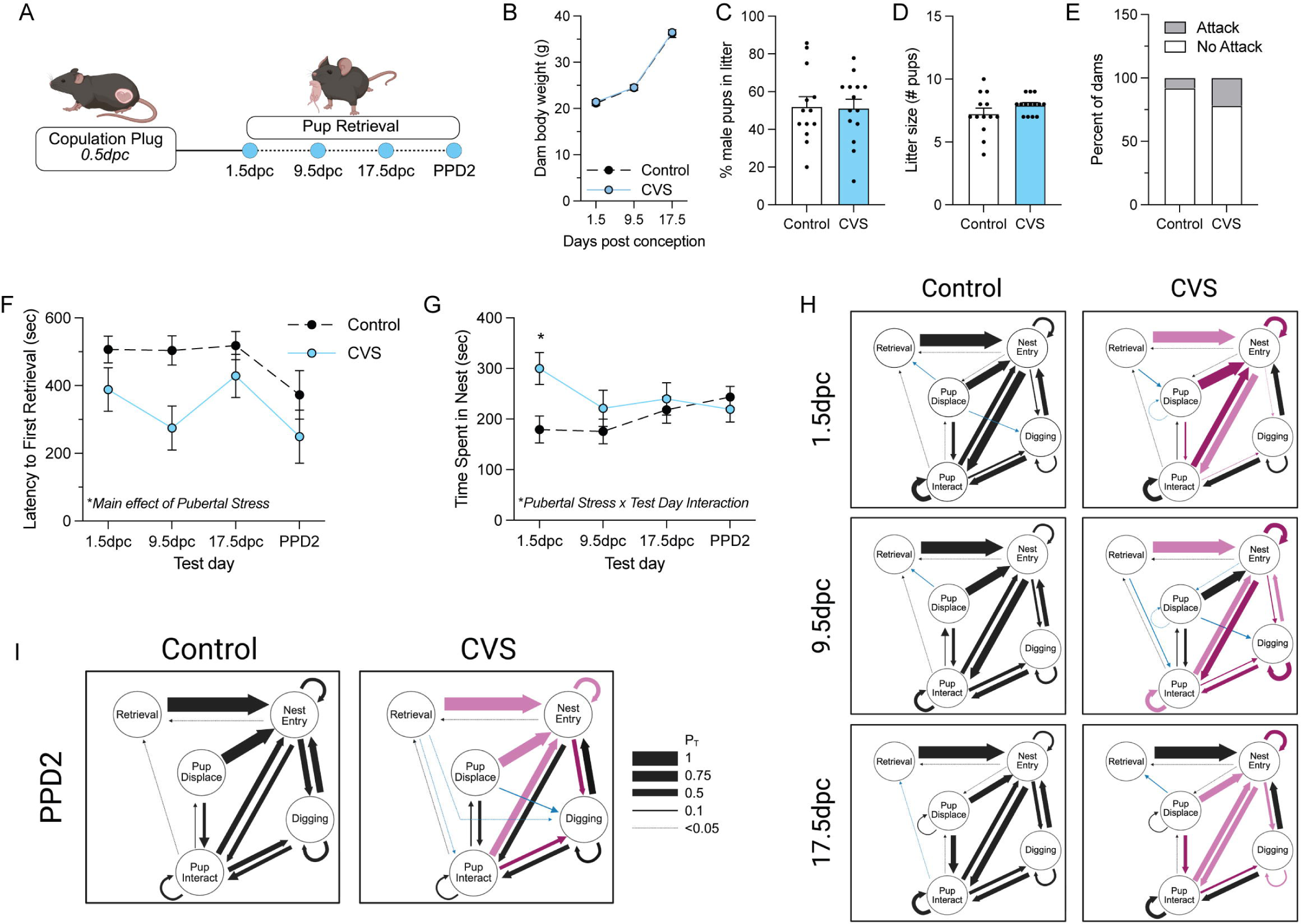
Pubertal stress led to changes in maternal behavior in adulthood during the pup retrieval task. **(A)** Females were bred in adulthood, and the presence of a copulation plug was considered 0.5dpc. Dams underwent a 10-minute pup retrieval task at 1.5dpc, 9.5dpc, 17.5dpc, and PPD2. **(B-D)** Pubertal stress did not alter general pregnancy outcomes, including (B) dam body weight throughout pregnancy, (C) sex ratio of offspring, and (D) litter size. **(E)** More pubertal CVS females (21.4%) were excluded during pup retrieval tasks for attacking pups than Controls (7.6%). In classical measures of outcomes in the pup retrieval task, pubertally stressed dams displayed alternations in behavior. Pubertal stress led to **(F)** decreased latency to retrieve the first pup and **(G)** significantly more time in the nest, but only at 1.5dpc. **(H,I)** More detailed analysis of behavior using Markov chain modeling shows that pubertal stress led to alterations of maternal behavior during pregnancy and postpartum. Arrows represent the transition probabilities (P_T_) that one behavior is followed by another behavior. Dark purple arrows represent transition probabilities that differed by >10% between Control and CVS, light purple arrows represent transition probabilities that differed between 5- 10%, and blue arrows represent transitions that were present in only one group. **(I)** Testing on PP2 revealed disorganized behavior in pubertally stressed females. CVS = chronic variable stress; dpc = days post conception; P_T_ = transition probability; PPD = postpartum day.

### Preconception pubertal stress led to a blunted corticosterone response to acute stress within and between testing days

We tested whether the previously observed blunted glucocorticoid response in CVS mice would be replicated using pup retrieval as a stressor (**Figure 3A**). There were no differences between the corticosterone response to pup retrieval stress at 1.5dpc, 9.5dpc, or 17.5dpc (**Figure 3B-3D**). There was an interaction between pubertal stress and test day on peak (30 min) corticosterone (*F_pubertal stress x test day_*(1.22, 28,02) = 4.106), *p* = 0.045), such that pubertally stressed mice displayed a lower corticosterone peak at 1.5dpc (*p* = 0.043; **Figure 3E**).

**Figure 3.**
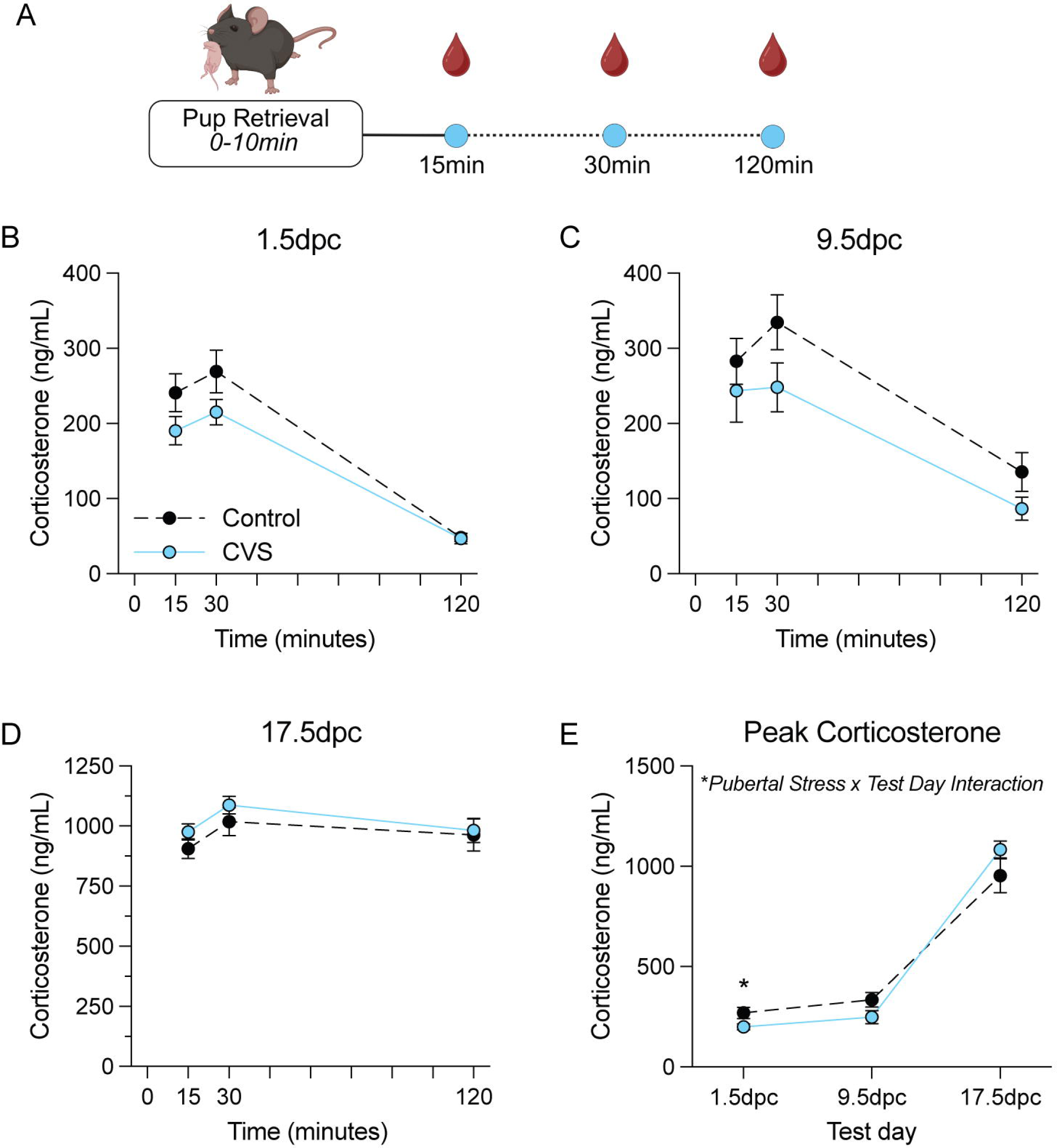
Pubertal stress led to a blunted HPA axis response at some time points during pregnancy. **(A)** Following the pup retrieval test, tail blood was collected at several time points to measure the corticosterone response to the test. **(B)** At 1.5dpc, there was a nonsignificant trend where pubertally stressed dams displayed a lower glucocorticoid response to acute stress. **(C)** A similar nonsignificant trend was observed at 9.5dpc. **(D)** There was no effect of pubertal stress on the glucocorticoid response to acute stress at 17.5dpc. **(E)** There was a significant interaction between pubertal stress and test day on peak 30-minute corticosterone, where pubertally stressed dams showed a blunted peak response at 1.5dpc. CVS = chronic variable stress; dpc = days post conception.

### Maternal preconception pubertal stress led to sex-specific changes in offspring physiology and behavior at different ages

One male and one female offspring per litter were retained for behavioral testing (**Figure 4A**). There was an interaction between age and dam pubertal stress on offspring body weight (**Figure 4B**, *F*(1,43) = 4.35, *p* = 0.045), such that adult offspring of CVS dams weighed less than adult offspring of Control dams (F(1,43) = 5.47, *p* = 0.024). Offspring underwent open field and social preference testing in adolescence and again in adulthood. We found a 3-way interaction between offspring age, dam pubertal stress, and sex on time in the center of the open field (**Figure 4C**, *F*(1,41) = 4.247, *p* = 0.046), such that male offspring of CVS dams spent more time in the center of the open field compared to each other groups in adolescence. There was a main effect of age on total distance traveled (*F*(1,41) = 113.3, *p* < 0.001), such that adult mice traveled less total distance than adolescents (**Figure 4D**). We found an effect of dam pubertal stress on social preference index during adolescence (*F*(1,40) = 4.842, *p* = 0.034), such that offspring of CVS dams displayed a greater social preference compared to offspring of Control dams (**Figure 4E**). There were no effects of dam pubertal stress or sex on social preference index during adulthood (**Figure 4F**).

**Figure 4.**
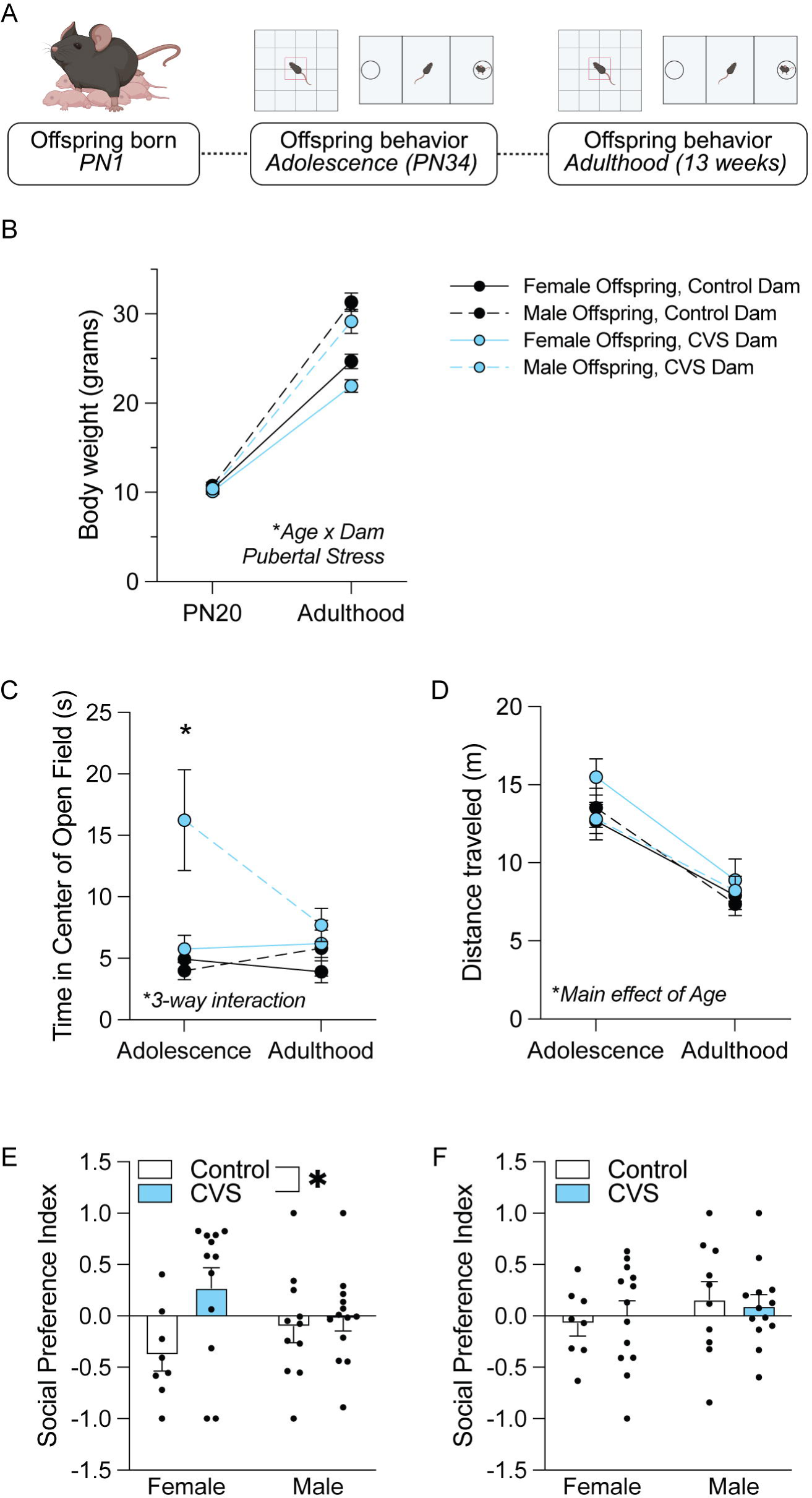
Maternal pubertal stress led to changes in offspring physiology and behavior at different time points. **(A)** Experimental timeline. Offspring of dams from Experiment 1 underwent a 5 minute open field test followed by a 10-minute social preference test during adolescence and adulthood. **(B)** There was a significant interaction between offspring age and dam pubertal stress, such that offspring of pubertally stressed dams weighed less than offspring of Control dams in adulthood, but not adolescence. **(C)** There was a significant interaction between dam pubertal stress, offspring sex, and offspring age, such that male offspring of pubertally stressed dams spent an increased time in the center of the open field compared to all other groups during adolescence, but not adulthood. **(D)** There was a main effect of offspring age on the distance travelled during the open field test such that offspring travelled a greater distance in adolescence compared to adulthood. **(E)** There was a main effect of pubertalchao stress on social preference index during adolescence such that offspring of pubertally stressed dams had a greater preference for the social zone. **(F)** There were no differences found on social preference index during adulthood. CVS = chronic variable stress. Dpc = days post conception. PN = postnatal day.

### Preconception pubertal stress led to a change in baseline expression of genes related to the expression of maternal behavior during late pregnancy

During late pregnancy, we examined the baseline gene expression of several genes established as necessary for the expression of maternal behavior (**Figure 5A**) [40–42,44]. Expression of *Pgr, Esr1,* and *Gal* were measured in the mPOA (**Figure 5B**). While we observed no group differences in the total number of mPOA cells detected (**Figure 5C**), other reported outcomes are expressed as a proportion of or average across detected cells. There were no differences in the expression *Esr1* or *Pgr* per cell (**Figure 5D**). Further, there were no differences in the percentage of all mPOA cells expressing *Esr1, Pgr,* or *Gal* (**Figure 5E**). There was an effect of pubertal stress on the percentage of *Gal*+ cells expressing only *Gal* (negative for *Esr1* or *Pgr*), where CVS mice displayed a greater percentage of cells without hormone receptor gene expression (**Figure 5F**). While there was no difference in the percent of *Gal+/Esr1+* cells, CVS females displayed a lower percentage of *Gal*+/*Pgr*+ cells. When all classification types were examined, it shows that CVS females generally had a lower percent of cells with *Pgr* expression (**Figure 5G**).

**Figure 5.**
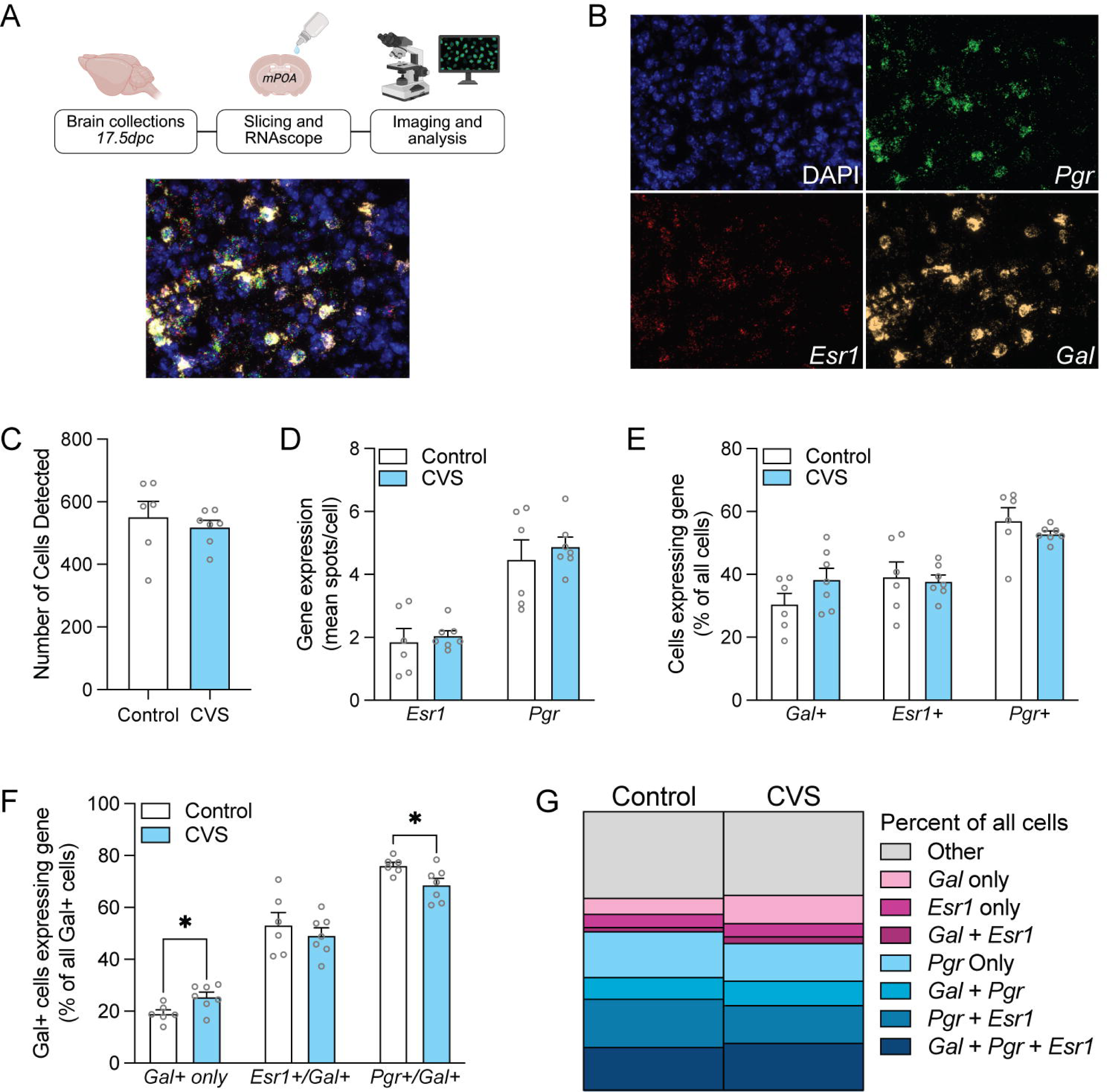
Pubertal stress led to changes in expression of maternal behavior relevant genes in the medial preoptic area of the hypothalamus. **(A)** Experimental timeline. The mPOA was collected at 17.5dpc for RNAscope *in situ* hybridization. Merged RNAscope image. **(B)** Visualization of separated DAPI, *Pgr, Esr1,* and *Gal* channels. **(C)** There was no effect of pubertal stress on the number of mPOA cells detected per slice. **(D)** There was no effect of pubertal stress on the mean amount of *Esr1* or *Pgr* expression detected per cell. **(E)** There was no significant effect of pubertal stress on the percentage of detected cells expressing *Gal, Esr1,* or *Pgr*. **(F)** There was a significant effect of pubertal stress on the percentage of *Gal-*expressing cells that did not express *Esr1* or *Pgr* such that pubertally stressed mice showed an increased percentage. There was no effect of pubertal stress on the percentage of *Gal-*expressing cells also expressing *Esr1.* There was a significant effect of pubertal stress on the percentage of *Gal-*expressing cells also expressing *Pgr* such that pubertally stressed mice showed a decreased percentage. **(G)** Stackplot representing the proportion of cells expressing each combination of genes of interest. CVS = chronic variable stress. Dpc = days post conception. MPOA = medial preoptic area.

## Discussion

Mood and anxiety disorders are highly prevalent in the perinatal window. While environmental factors, such as early life stress, are known risk factors for such disorders, little is known about how the early life environment interacts with later pregnancy to further exacerbate this risk. To address such questions, we utilize a translationally relevant mouse model to investigate how early life stress leads to long-term reprogramming in the brain, lifespan-relevant behavior, and physiology that is revealed by pregnancy.

Our previous work established that in both peripartum humans and mice, stress during puberty led to a blunted glucocorticoid response to acute stress. Here, to better understand whether this phenotype has implications for behavior, we performed a pup retrieval task repeatedly across pregnancy and early postpartum. Interestingly, pubertal stress led to faster pup retrieval across pregnancy and postpartum. However, it is important to note that faster retrieval may not always be indicative of better maternal care. Other studies have found that, while pup-directed behaviors may be increased following a manipulation, the pattern or sequence of maternal care changes in a way that may be maladaptive to offspring [55–58]. To better understand the pattern of maternal behavior, we used a Markov Chain Model to produce transition probabilities for each pairing of scored behaviors. These results revealed a pattern where pubertally stressed females were more likely to repeat the same behavior multiple times in a row compared to controls. This is most noticeable in the increased probability of repeated nest entry on all test days. This is consistent with findings from other models of maternal stress, which have identified an increased frequency of nest exit and entry [55]. In addition, we found that pubertally stressed dams spent a significantly greater amount of time in the nest only on the first day of testing. Finally, pubertally stressed dams were more than twice as likely to attack pups and be removed from testing (22%) than nonstressed dams (8%). Taken together, these findings suggest that while pubertally stressed dams retrieved more quickly throughout pregnancy and postpartum, this may not necessarily translate to an increase in the amount of time spent caring for pups after retrieval.

Immediately following the pup retrieval task, blood was collected at several timepoints to examine the plasma corticosterone response. Previously, we found that pubertal stress led to a blunted stress response to an acute restraint stress in late pregnancy [21,31]. Here, we observed a similar trend in early- and mid- pregnancy, where pubertally stressed dams showed a slightly decreased corticosterone response. Pubertally stressed mice displayed a significantly lower peak corticosterone response at 1.5dpc, but not on other testing days. At 17.5 days post conception, the same timepoint we previously observed a blunted response following restraint stress [21,31], we saw no difference in the corticosterone response to acute stress. Several factors may have contributed to these differences from our previous findings. First, the previously established blunted corticosterone response was in response to an acute restraint stress. Here, exposure to the retrieval test and the presence of novel pups in the home cage acted as the stressors. Although both are effective in eliciting a corticosterone response, not all stressors do so through the same neurobiological pathways. In fact, while there is some overlap in the pattern of activation, physical restraint stress and social stress are known to elicit distinct patterns of hypothalamic activation [59]. The prior experience of pubertal stress may program lasting changes in the response to some stressful stimuli, but not others. Second, the use of repeated measures testing limits comparisons to our past work. Unlike our previous work, mice in the current study were exposed to the pup retrieval task three times throughout pregnancy. Repeated exposures to the same stimulus can result in habituation of the HPA axis response over time [60,61]. The repeated measures design used has several strengths, including that it reduces the number of mice required and allows for observation of behavior change over time within the same animal. However, one limitation is that it is unknown whether changes in stress reactivity over time are a result of habituation to the stressor. This is further complicated by the fact that the HPA axis response is known to change throughout pregnancy. Further work is needed to understand whether these changes are due to the nature of the acute stressor or repeated testing. Nonetheless, these results confirm that pubertal stress programs lasting changes in HPA axis reactivity.

Using open field and social preference tasks, we assessed the behavior of male and female offspring of the dams that underwent the pup retrieval task. These tasks were administered during adolescence and adulthood in the same mice. The open field task is sometimes conceptualized as an approach-avoidance conflict task, such that it pits the rodent’s tendency to explore its environment against its tendency to remain in closed or protected areas [62–65]. Here, we found that male offspring of pubertally stressed dams spent more time in the center of the open field compared to all other groups during adolescence. In a natural setting, spending too much time in open and exposed areas may unnecessarily expose rodents to predators. Therefore, an increase in the amount of time spent in the center compared to controls could be interpreted as increased risk-taking behavior [66,67]. These data fit interestingly into existing work, which has found that males are generally more prone to risk-taking behavior, particularly during adolescence or after experiencing a stressor [68–71]. Here, the effect of maternal preconception stress is no longer present in adult offspring. This aligns with findings in human subjects, where risk-taking behaviors peak in adolescence and begin to decline throughout young adulthood [72]. Our data support findings that suggest adolescence is a particularly sensitive window for risk-taking behaviors, and that maternal preconceptions exposures can intergenerationally increase such risk.

Interestingly, several human studies have revealed that while males are more prone to risk-taking compared to females in several domains, differences in social or relational risk-taking are negligible [73,74]. A functional imaging study in humans found that stimuli associated with either physical or social risk led to distinct patterns of neural activation [75]. Here, we found that offspring of pubertally stressed dams showed an increase in social preference during adolescence, but not adulthood. While not significant, it is worth noting that this effect is largely driven by female offspring of pubertally stressed dams. Taken together with our findings from the open field task, our data suggest that dam pubertal stress may lead to sex- and domain- specific increases in risk-taking behavior during the sensitive period of adolescence. Future work will be needed to define when this window of risk closes, as our data show it is closed by adulthood. Nonetheless, this means that maternal preconception stress during puberty may lead to an increased window of behavioral vulnerability in offspring during adolescence.

While we identified no effects of maternal pubertal stress in offspring behavior during adulthood, we did find that offspring of CVS dams weighed less compared to those of Control dams. This finding indicates that some effects of maternal pubertal stress last well into the lifespan of offspring and is consistent with other work showing longer duration preconception stress is associated with altered metabolic outcomes in female offspring [76]. These differences in the behavior and physiology of the offspring of dams that underwent pubertal stress further strengthen the conclusion that some aspects of maternal physiology or behavior may be maladaptive and contribute to lasting changes in offspring.

Finally, we examined the expression of several genes important for maternal behavior (*Gal, Pgr, Esr1*) in the medial preoptic area of the hypothalamus. Progesterone and estrogen signaling have long been established to play an important role in the expression of maternal behavior [44,77–79]. In mammals, circulating progesterone levels gradually increase throughout pregnancy and sharply drop following parturition [80–82]. Importantly, progesterone is known to down-regulate ERα and ERβ receptors [83,84]. Following parturition, the sharp decrease in progesterone levels therefore leads to an increased sensitivity to estrogen signaling [79], which is thought to contribute to the initiation of maternal behavior in mammals [85]. In rats, estrogen administration has been shown to stimulate behavior most potently when it is administered after termination of pregnancy [35]. Further, administration of progesterone blocks the stimulation of maternal behavior by estrogen [35]. We found that while *Esr1* expression generally did not vary due to pubertal stress, the percentage of *Gal*-positive cells expressing *Pgr* was decreased in pubertally stressed mice. Ammari et al. (2023) found that ablation of mPOA galanin neurons expressing either *Esr1* or *Pgr* was enough to impair maternal behavior to the same degree that was observed following an mPOA- wide knockout of these receptors [44]. Our findings suggest that pubertal stress may attenuate the peak effects of progesterone signaling observed in late pregnancy, which, in turn, may lead to lesser progesterone withdrawal following parturition. Further work is needed to map the trajectories of progesterone and estrogen binding throughout the peripartum window following pubertal stress. However, these data suggest that the differences in maternal behavior we find here may be related to changes in progesterone activity on galanin-expressing neurons in the mPOA.

In conclusion, we demonstrate that puberty is a sensitive window for stress to influence later behavior, physiology, and neurobiology during the peripartum window. An intergenerational effect of stress was evident, as offspring of moms stressed during preconception had altered behavioral and metabolic outcomes. These findings expand upon our prior work in mice demonstrating disruptions to endophenotypes of neuropsychiatric disorders, stress physiology, and the epigenetic landscape in the hypothalamus, and mirrors what we find in human mothers and offspring [20,21,30,32]. Moreover, these findings provide additional insight into the risk factors, even those that occur long before pregnancy, for disrupted outcomes for both mom and offspring.

## Author Contributions

MMN: Substantial contributions to the acquisition, analysis, or interpretation of data for the work, drafting the work or revising it critically for important intellectual content

LAML: Substantial contributions to the acquisition of the work

IML: Substantial contributions to the acquisition of the work

MAF: Substantial contributions to the acquisition of the work

KYM: Substantial contributions to the acquisition of the work

KEM: Substantial contributions to the conception or design of the work; or the acquisition, analysis, or interpretation of data for the work; and drafting the work or revising it critically for important intellectual content; and final approval of the version to be published. Agreement to be accountable for all aspects of the work in ensuring that questions related to the accuracy or integrity of any part of the work are appropriately investigated and resolved.

