## Supplemental Materials for "Preconception stress in mice alters maternal behavioral, neural, and endocrine outcomes, and produces sex-specific disruptions in offspring"

*Mice*

Mice used were in-house, heterozygous Crh-IRES-Cre:Ai14, expressing tdTomato in CRF neurons on a mixed C57BL/6 x 129S1/SvImJ background or the offspring of such mice (Experiment 1, N=27; Experiment 2, N=48; Experiment 3, N=13). Mice were housed in a 12:12 hour light:dark cycle with access to food and water *ad libitum*. All procedures were approved by the West Virginia University Institutional Animal Care and Use Committee and were conducted in accordance with the National Institutes of Health Guide for the Care and Use of Laboratory Animals.

*Chronic Variable Stress*

Female mice were assigned to the Control or CVS condition, balancing for litter of origin and bodyweight at postnatal day (PN) 20. Mice assigned to the CVS group underwent two weeks of daily stressors targeting a variety of sensory modalities beginning on PN21 and ending on PN34, inclusive. Once a day, two stressors from the following list were administered simultaneously: auditory stressors (owl screech, white noise), olfactory stressors (predator odor, ethanol odor), or tactile stressors (restraint in a conical tube, wet bedding, multiple cage change, exposure to novel stimulus, no bedding). Descriptions of each stressor are in Supplemental Table S1. All stressors were administered for the first two hours of the light cycle, with the exception of restraint stress, which had a duration of 15 minutes. CVS mice were singly housed at PN21 and for the duration of CVS, but were pair housed with same-sex, same-condition littermates at PN34 following the conclusion of CVS. Control mice were weaned at PN28 and pair-housed with same-sex littermates as previously.

*Pup Retrieval*

Pregnant female mice underwent a home cage pup retrieval task at four different testing days, at 1.5 dpc, 9.5 dpc, 17.5 dpc, and postpartum day (PPD) 2. Testing began one hour after the start of the light cycle. Because female mice were pregnant during some testing days, probe pups (aged PN2-6) from a donor dam were required for the retrieval task. One pup was placed in the corners far from the nest in the home cage of the test female, with a male always placed on the left corner and a female on the right corner. In the case of PPD2 females, the entire litter was removed during the test, except the two pups used for retrieval. The test female was then placed in the nest and her behavior was recorded for 10 min. If mice acted aggressively toward pups, the test was ended early, and they were excluded from analysis of pup retrieval. Aggression was defined as the adult mouse biting, lunging at, or grooming excessively to a point that elicited audible cries from the pup.

Behavior was hand-scored in Anymaze. For a full list of scored behaviors and definitions used, see **Table S2**. Behaviors were analyzed in bouts. If the same behavior, for example digging, was paused and resumed with less than a one second pause, and no other scored behavior occurred in that window, these occurrences were considered one bout of digging. If the pause was greater than one second or another scored behavior occurred during that window, then it was treated as a new bout.

To produce probe pups for the pup retrieval test, a separate cohort of female C57BL/6 x 129S1/SvImJ mice were paired with experienced male breeders. Females were bred for four nights, 22-25 days prior to the start of each round of behavioral testing. After breeding, they were singly housed and remained undisturbed until PN2. Probe pups were used for no more than 2 tests each day for no more than 2 total days of testing. Prior to testing, probe pups were removed from their dam cage and placed in a cage on a heating pad in the testing room. Prior to each test, the nest of the female was placed in the back right corner of the cage to ensure a consistent location across mice.

*Glucocorticoid response to acute stress*

On 1.5 dpc, 9.5 dpc, and 17.5 dpc testing days, 10 ul of blood was collected from females from a small nick at the end of the tail at 15 minutes, 30 minutes, and 120 minutes following the start of pup retrieval testing. Blood samples were immediately mixed with 5 µl of 50 mM EDTA and kept on ice until centrifugation for 10 min at 5,000 rpm at 4°C. Plasma was collected and stored at -80°C until analysis. Corticosterone concentration was determined from blood plasma by 125I-corticosterone radioimmunoassay (MP Biomedicals), per manufacturer instructions.

*Open Field*

Offspring were placed into the center of a 40 x 40 centimeter apparatus and were allowed to freely explore for 5 minutes. Tracking of location and distance travelled was performed by AnyMaze. The center zone was defined as the centermost 100 square centimeters. The corner zone was defined as 100 square centimeters in any of the four corners of the apparatus.

*Social Preference*

A 60 x 40 centimeter 3-chamber box was used for social preference testing, where each chamber was 20 x 40 centimeters. The two innermost walls separating the chambers each contained a 7 x 7 centimeter opening through which the mouse could cross between chambers. Mice were allowed to habituate to the apparatus, which included two empty cups placed at the center of the far zones of the chamber, for 5 minutes. For the test, a same-sex, same-age novel probe mouse was placed into the cup in one of the far zones. To account for side preference, the probe mouse was placed in the side where the test mouse spent the least amount of time during habituation. Then, the test mouse was returned to the chamber and allowed to explore for 5 minutes. Tracking of location and distance traveled was performed by AnyMaze. To calculate the social preference index, the following formula was used: (time in social zone - time in nonsocial zone)/(time in social zone + time in nonsocial zone).

*Tissue collection*

At baseline on the morning of 17.5 dpc, mice were anesthetized via 5% isoflurane, then euthanized via decapitation. Brains were extracted, immediately placed on dry ice, and stored at -80°C. Brains were cryosectioned at -20°C and the medial preoptic area (mPOA) was collected (0.25mm through -0.23mm relative to bregma). 4, 10um slices were collected in a set of three slides. Each slide in the set represented the full anterior to posterior range for the mPOA.

*In situ hybridization*

Gene expression analyses were performed using the RNAscope Fluorescent Multiplex Reagent Kit (version 2, ACDBio) according to manufacturer instructions for fresh frozen tissue (RNAscopeTM Multiplex Fluorescent Reagent Kit v2 User Manual, 2022). Protease IV digestion time was reduced from 30 minutes to 15 minutes to correct for overdigestion observed when piloting the assay. Probes targeted *Gal* (RNAscope Probe- Mm-Gal), *Pgr* (RNAscope Probe- Mm-Pgr-C2), and *Esr1* (RNAscope Probe- Mm-Esr1-C3). Probe channel, fluorophore, and fluorophore dilution were as follows: *Gal* (Channel 1, TSA Vivid 570, 1:3000); *Pgr* (Channel 2, TSA Vivid 520, 1:1500), *Esr1* (Channel 3, TSA Vivid 650, 1:3000).

*In situ hybridization imaging and quantification*

Fluorescent images were acquired using a 40x objective on an Olympus BX43 Upright Microscope equipped with X-Cite 120LED Boost High Power Illumination System. The following LED filter cubes, exposure time, and X-Cite 120LED Boost light intensity were used for each fluorophore: TSA Vivid 570 (RFP, 3sec, 1% intensity); TSA Vivid 520 (GFP, 10sec, 2% intensity); TSA Vivid 650 (Cy5, 3sec, 1% intensity). Acquisition settings were determined based on piloting using RNAscope positive control probe. Distance of the center of the ROI relative to the base of the third ventricle was used to capture region of interest. Measurement of the distance were determined based on the Allen Mouse Brain Atlas as follows: Slice 1 (250 μm lateral, 100 μm dorsal); Slices 2,3 (400 μm lateral, 100 μm dorsal); Slice 4 (500 μm lateral, 250 μm dorsal).

Images were exported as grayscale 8-bit files and merged as converted to 4-channel hyperstacks in ImageJ. Images were uploaded to QuPath for cell and spot identification. The positive cellular detection tool was used and a cell radius of 15μm, including nuclei, was used, and cells were labeled as either positive or negative for *Gal*. Subcellular detection was performed for *Pgr* and *Esr1*. Positive threshold values were determined based on piloting using RNAscope positive control probes. The measurements reported (e.g., mean spots/cell, percent of positive cells) were determined based on the Guide for RNAscope Data Analysis, as published by the manufacturer.

**Statistical Analyses**

For duration measures in the pup retrieval task, a 2 x 4 ANOVA was used to test main effects of pubertal stress and test day, as well as the interaction between these factors. A 2 x 3 ANOVA was used to test effects and interactions of pubertal stress and time (repeated measure) on plasma corticosterone and to test effects and interactions of pubertal stress and test day on 30-min peak plasma corticosterone. For open field behavior measures, a 2 x 2 x 2 ANOVA was used to test the effects and interactions of sex, dam pubertal stress, and age of offspring (adolescent vs adult). For social preference behavior measures, a 2 x 2 ANOVA was used to test the effects of sex and dam pubertal stress within each testing age. For all ANOVAs, if the assumption of sphericity was violated, the Geisser-Greenhouse correction was used. A Markov Chain analysis was performed on the transition probabilities of maternal behavior during pup retrieval. Independent sample t-tests (Control vs CVS) were used to compare gene expression and cell population measures for *in situ* hybridization. For all measures, the ROUT method was used to identify outliers. P value set to 0.05. Full statistical output is presented in **Tables S3-S9**.

### Supplemental Tables

**Table S1.** Full description of stressors.

| | |
|---|---|
| **Tactile stressors** | |
| Wet bedding | Bedding and nest are made completely wet with tap water. Enough water is used so that bedding and nest are fully saturated but there is no standing water in the cage. |
| Restraint | Unlike other stressors, which have a duration of 2 hours, restraint stress lasts for only 15 minutes. Mice are placed in a 50 ml conical tube that has been modified by adding holes in their home cage. To make the adult restraint tubes function properly for pubertal mice, tubes are prepared with either four (for mice age PN21-27) or three (for mice age PN28-34) marbles in the distal portion of the tube. This prevents the small pubertal mice from turning around in the tube. |
| Wire mesh | A sheet of wire mesh is placed on top of the bedding and nest of the home cage so that the mouse cannot access them. The mesh remains for the duration of stress. |
| Multiple cage change | The home cage is changed three times in the duration of two hours- at the beginning of the two-hour stress period, halfway through the two-hour period, and at the conclusion of stress. Only the cage bottom (containing the corncob bedding) is changed. The other elements of the cage, including the lid, water bottle, and wire top remain the same. At the conclusion of this stressor, one of the cage bottoms is retained to be used for no bedding. |
| No bedding | All bedding and the nest is removed from the home cage for the duration of stress. |
| Novel stimulus | Marbles are introduced into the home cage to act as a novel stimulus. Eight marbles are placed as evenly as possible in a 2x4 array, and are placed directly atop the nest when necessary. The marbles remain for the duration of stress. |
| **Auditory stressors** | |
| White noise | White noise is played from a speaker in the room where mice undergo stress for the duration of stress. |
| Owl screech | Owl screeches, obtained from the Cornell Lab of Ornithology, are played from a speaker in the room where mice undergo stress for the duration of stress. |
| **Olfactory stressors** | |
| Ethanol odor | A 15 mL conical tube with approximately 5 ml of 70% ethanol is placed in the food hopper for the duration of stress. |
| Predator odor | A 15 mL conical tube with approximately 5 ml of Puma odor - 2-Phenethylamine (CAS 64-04-0), 1:200 in mineral oil (Sigma Aldrich M5904) is placed in the food hopper for the duration of stress. |

**Table S2.** Pup retrieval task behavioral scoring definitions

| Behavior | Definition | Key Function |
|---|---|---|
| Contact Latency | The time of first contact of the female's nose with either of the pups. | Point |
| Digging | Start when the female begins digging in the corncob bedding with her nose or paws. Stop when she ceases. Duration will be the total amount of time the female spends doing so. | State |
| Nest Entry | Start when the female enters the nest with all four paws. Stop when she exits the nest with at least one of her paws stepping out of the nest. Poking only her head out of the nest does not count as an exit. Duration will be defined as the total amount of time the female spends in the nest. | State |
| Pup Displacement | The moment a pup has been put down after being carried by the female in her mouth to a new location, but part or all of the pup's body remains outside the nest | Point |
| Pup Interaction | Start when the female interacts closely with the pup(s) using her nose, mouth, or paws. This includes carrying the pup. Stop when she ceases contact with the pup(s). Duration will be defined as the total amount of time she spends doing so. | State |
| Retrieval Latency | The moment a pup has been carried/pulled completely into the nest by the dam using her mouth, and no part of the pup's body remains outside the nest. Not scored if a pup was covered with nesting material being manipulated by the dam, or if a pup moved itself into the nest. | Point |
| One Pup in Nest | The total duration of time one of two pups is fully inside the nest. | State |
| Two Pups in Nest | The total duration of time both pups are fully inside the nest. | State |

**Table S3.** Full statistical reporting from independent t-test analyses in Experiment 1

| Outcome | t ratio | df | P value | Significance |
|---|---|---|---|---|
| % Male Pups in Litter *(Figure 2C)* | 0.101 | 25 | 0.92 | ns |
| Litter Size *(Figure 2D)* | 1.446 | 25 | 0.161 | ns |

**Table S4.** Full statistical reporting from 2-way ANOVA analyses in Experiment 1

| Outcome | Factor | F value | P value | Significance |
|---|---|---|---|---|
| *Dam body weight (Figure 2B)* | | | | |
| | Test Day | F(1.251, 31.29) = 1230 | <0.001 | * |
| | Pubertal Stress | F(1, 25) = 0.08 | 0.779 | ns |
| | Test Day x Pubertal Stress | F(1.251, 31.29) = 0.057 | 0.864 | ns |
| *Latency to First Retrieval (Figure 2F)* | | | | |
| | Test Day | F(2.212, 46.45) = 5.613 | 0.005 | * |
| | Pubertal Stress | F(1, 21) = 4.819 | 0.04 | * |
| | Test Day x Pubertal Stress | F(2.212, 46.45) = 0.978 | 0.391 | ns |
| *Time Spent in Nest (Figure 2G)* | | | | |
| | Test Day | F(2.566, 53.89) = 1.537 | 0.22 | ns |
| | Pubertal Stress | F(1,21) = 2.402 | 0.136 | ns |
| | Test Day x Pubertal Stress | F(2.566, 53.89) = 2.959 | 0.048 | * |
| *1.5dpc Corticosterone (Figure 3B)* | | | | |
| | Time | F(1.583, 39.57) = 129.8 | <0.001 | * |
| | Pubertal Stress | F(1, 25) = 2.686 | 0.114 | ns |
| | Test Day x Pubertal Stress | F(1.583, 39.57) = 2.503 | 0.106 | ns |
| *9.5dpc Corticosterone (Figure 3C)* | | | | |
| | Time | F(1.442, 33.16) = 49.27 | <.001 | * |
| | Pubertal Stress | F(1, 23) = 2.306 | 0.143 | ns |
| | Test Day x Pubertal Stress | F(1.442, 33.16) = 0.815 | 0.415 | ns |
| *17.5dpc Corticosterone (Figure 3D)* | | | | |
| | Time | F(1.376, 33.03) = 6.356 | 0.01 | * |
| | Pubertal Stress | F(1, 24) = 0.856 | 0.364 | ns |
| | Test Day x Pubertal Stress | F(1.376, 33.03) = 0.409 | 0.592 | ns |
| *Peak Corticosterone (Figure 3E)* | | | | |
| | Test Day | F(1.218, 28.02) = 213.9 | <0.001 | * |
| | Pubertal Stress | F(1, 23) = 0.036 | 0.852 | ns |
| | Test Day x Pubertal Stress | F(1.218, 28.02) = 4.106 | 0.045 | * |

*Testing Day (1.5dpc, 9.5dpc, 17.5dpc, PPD2); Pubertal Stress (Control, CVS).*

**Table S5.** Full statistical reporting from Tukey's multiple comparisons in Experiment 1.

| Outcome | Mean difference | 95% CI | | P value | Significance |
|---|---|---|---|---|---|
| | | Lower | Upper | | |
| *Time Spent in Nest (Figure 2G)* | | | | | |
| 1.5dpc, Control - CVS | -120.6 | -206.8 | -34.42 | 0.009 | * |
| 9.5dpc, Control - CVS | -45.79 | -136.5 | 44.9 | 0.303 | ns |
| 17.5dpc, Control - CVS | 21.61 | -107.9 | 64.64 | 0.607 | ns |
| *Peak corticosterone (Figure 3E)* | | | | | |
| 1.5dpc, Control - CVS | 69.53 | 2.337 | 136.7 | 0.043 | * |
| 9.5dpc, Control - CVS | 86.54 | -14.54 | 187.6 | 0.09 | ns |
| 17.5dpc, Control - CVS | -131.2 | -330.5 | 68.2 | 0.184 | ns |

**Table S6.** Full statistical reporting from ANOVA analyses in Experiment 2

| Outcome | Factor | F value | P value | Significance |
|---|---|---|---|---|
| *Offspring Body weight (Figure 4B)* | | | | |
| | Age | F(1, 43) = 970 | <0.001 | * |
| | Dam Pubertal Stress | F(1, 43) = 5.356 | 0.026 | * |
| | Sex | F(1, 43) = 37.32 | <0.001 | * |
| | Age x Dam Pubertal Stress | F(1, 43) = 4.254 | 0.045 | * |
| | Age x Sex | F(1, 43) = 39.42 | <0.001 | * |
| | Dam Pubertal Stress x Sex | F(1, 43) = 0.048 | 0.828 | ns |
| | Age x Dam Pubertal Stress x Sex | F(1, 43) = 0.104 | 0.749 | ns |
| *Offspring Body Weight – 2-way ANOVA at PN21 (Figure 4B)* | | | | |
| | Sex | F(1,43) = 0.749 | 0.392 | ns |
| | Dam Pubertal Stress | F(1,43) = 0.565 | 0.456 | ns |
| | Dam Pubertal Stress x Sex | F(1,43) = 0.010 | 0.921 | ns |
| *Offspring Body Weight – 2-way ANOVA in adulthood (Figure 4B)* | | | | |
| | Sex | F(1,43) = 43.42 | <0.001 | * |
| | Dam Pubertal Stress | F(1,43) = 5.47 | 0.024 | * |
| | Dam Pubertal Stress x Sex | F(1,43) = 0.081 | 0.777 | ns |
| *Time in Center of Open Field (Figure 4C)* | | | | |
| | Age | F(1, 41) = 1.604 | 0.213 | ns |
| | Dam Pubertal Stress | F(1, 41) = 8.152 | 0.007 | * |
| | Sex | F(1, 41) = 4.602 | 0.038 | * |
| | Age x Dam Pubertal Stress | F(1, 41) = 2.417 | 0.128 | ns |
| | Age x Sex | F(1, 41) = 1.124 | 0.295 | ns |
| | Dam Pubertal Stress x Sex | F(1, 41) = 3.333 | 0.075 | ns |
| | Age x Dam Pubertal Stress x Sex | F(1, 41) = 4.247 | 0.046 | * |
| *Time in Center of Open Field – 2-way ANOVA in adolescence (Figure 4C)* | | | | |
| | Sex | F(1,41) = 3.504 | 0.068 | ns |
| | Dam Pubertal Stress | F(1,41) = 6.617 | 0.014 | * |
| | Dam Pubertal Stress x Sex | F(1,41) = 5.048 | 0.030 | * |
| *Time in Center of Open Field – 2-way ANOVA in adulthood (Figure 4C)* | | | | |
| | Sex | F(1,41) = 1.333 | 0.255 | ns |
| | Dam Pubertal Stress | F(1,41) = 1.955 | 0.170 | ns |
| | Dam Pubertal Stress x Sex | F(1,41) = 0.018 | 0.893 | ns |
| *Distance traveled (Figure 4D)* | | | | |
| | Age | F(1, 41) = 113.3 | <0.001 | * |
| | Dam Pubertal Stress | F(1, 41) = 1.009 | 0.321 | ns |

| | | | |
|---|---|---|---|
| Sex | F(1, 41) = 0.607 | 0.441 | ns |
| Age x Dam Pubertal Stress | F(1, 41) = 0.018 | 0.895 | ns |
| Age x Sex | F(1, 41) = 0.084 | 0.774 | ns |
| Dam Pubertal Stress x Sex | F(1, 41) = 0.856 | 0.36 | ns |
| Age x Dam Pubertal Stress x Sex | F(1, 41) = 2.726 | 0.106 | ns |
| *Social preference index (adolescence) (Figure 4E)* | | | |
| Dam Pubertal Stress | F(1, 40) = 4.842 | 0.034 | * |
| Sex | F(1, 40) = 0.003 | 0.954 | ns |
| Dam Pubertal Stress x Sex | F(1, 40) = 2.973 | 0.092 | ns |
| *Social preference index (adulthood) (Figure 4F)* | | | |
| Dam Pubertal Stress | F(1, 40) = 0.004 | 0.953 | ns |
| Sex | F(1, 40) = 1.279 | 0.265 | ns |
| Dam Pubertal Stress x Sex | F(1, 40) = 0.247 | 0.623 | ns |

*Age (Adolescence, Adulthood); Maternal Stress (Maternal Control, Maternal CVS); Sex (Male, Female)*

**Table S7.** Full statistical reporting from Tukey's multiple comparisons in Experiment 2.

| Outcome | Mean difference | 95% CI | | P value | Significance |
|---|---|---|---|---|---|
| | | Lower | Upper | | |
| *Time in Center of Open Field – 2-way ANOVA in adolescence (Figure 4C)* | | | | | |
| Female, Control - Female, CVS | -0.828 | -10.61 | 8.95 | 0.996 | ns |
| Female, Control - Male, Control | 0.953 | -9.407 | 11.31 | 0.995 | ns |
| Female, Control - Male, CVS | -11.31 | -21.08 | -1.527 | 0.018 | * |
| Female, CVS - Male, Control | 1.782 | -7.703 | 11.27 | 0.958 | ns |
| Female, CVS - Male, CVS | -10.48 | -19.32 | -1.632 | 0.015 | * |
| Male, Control - Male, CVS | -12.26 | -21.74 | -2.774 | 0.007 | * |

**Table S8.** Full statistical reporting from independent t-test analyses in Experiment 3

| Outcome | t ratio | df | P value | Significance |
|---|---|---|---|---|
| Number of Cells Detected *(Figure 5C)* | 0.649 | 11 | 0.53 | ns |
| Esr1 (mean spots/cell) *(Figure 5D)* | 0.45 | 11 | 0.661 | ns |
| Pgr (mean spots/cell) *(Figure 5D)* | 0.604 | 11 | 0.558 | ns |
| Gal+ cells (% all cells) *(Figure 5E)* | 1.569 | 11 | 0.145 | ns |
| Esr1+ cells (% all cells) *(Figure 5E)* | 0.285 | 11 | 0.781 | ns |
| Pgr+ cells (% all cells) *(Figure 5E)* | 1.07 | 11 | 0.308 | ns |
| Gal+ only cells (% all Gal+ cells) *(Figure 5F)* | 2.708 | 11 | 0.02 | * |
| Esr1+/Gal+ cells (% all Gal+ cells) *(Figure 5F)* | 0.752 | 11 | 0.468 | ns |
| Pgr+/Gal+ cells (% all Gal+ cells) *(Figure 5F)* | 2.535 | 11 | 0.023 | * |
